## Supplementary Figures for "RNA G-quadruplex dynamic steers the crosstalk between protein synthesis and energy metabolism"

Dumas, *et al.*, Supplementary Fig 1

**a**

| RNA G-quadruplexes | | GO term | Annotated | Significant | pv.fisher.BH |
|---|---|---|---|---|---|
| Experimentally identified RG4-seq, *in vitro* | *Kwok_2016* (K+; PDS) | Mitochondrion | 1570 | 550 | 2.16E-15 |
| | | Mitochondrial part | 1023 | 344 | 1.96E-07 |
| | | Mitochondrial outer membrane | 180 | 74 | 6.44E-05 |
| | | Mitochondrial matrix | 462 | 161 | 1.04E-04 |
| | *Guo_2016* (K+) | Mitochondrion | 1570 | 463 | 2.04E-09 |
| | | Mitochondrial outer membrane | 180 | 67 | 8.19E-05 |
| Predicted Algorithm, GQ score > 21 | | Mitochondrion | 1570 | 1090 | 6.64E-06 |
| | | Mitochondrial outer membrane | 180 | 140 | 2.65E-04 |

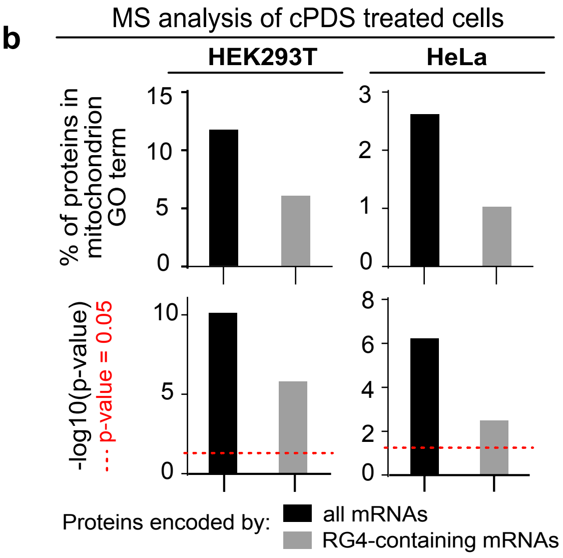


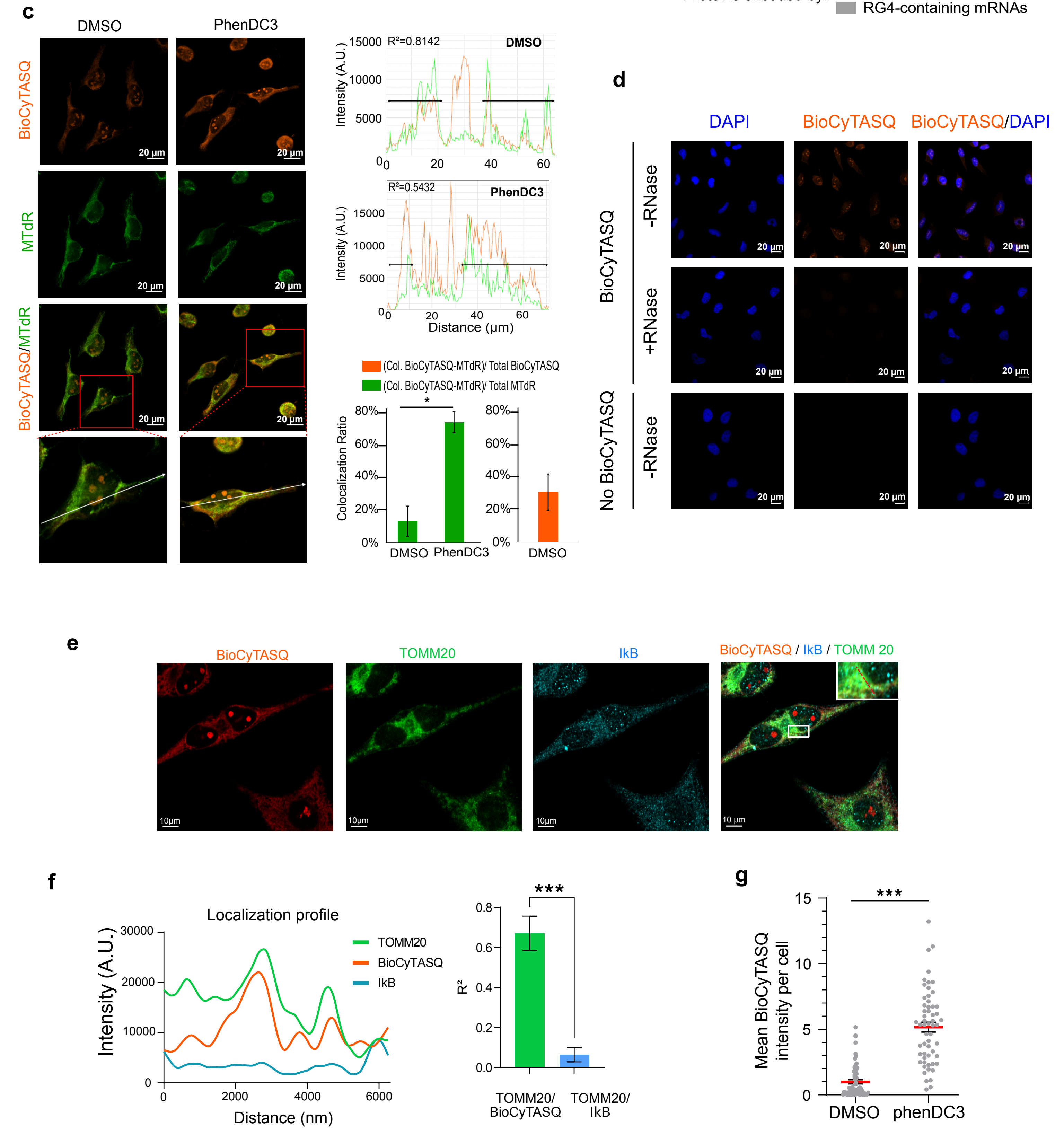

**Supplementary Figure 1 (related to Figure 1). RG4 localization in mitochondrial mRNAs (a)** Gene ontology enrichment analysis of RG4-containing mRNAs either experimentally validated based on [1,2] or predicted based on [3]. **(b)** Gene ontology enrichment analysis, along with the -log10 p-value of the enrichment, of proteins encoded by mRNAs containing predicted RG4s and identified by mass spectrometry analysis to be differentially expressed after 20 µM carboxypyridostatin (cPDS) treatment for 48 h in HEK293T or HeLa cells (n=5 independent experiments). **(c)** RG4 detection in live cells using the BioCyTASQ probe (1 µM for 24 h) and mitochondria labelling using MitoTracker Deep Red (MTdR) of HeLa cells treated with DMSO or 20 µM of PhenDC3 for 24 h. Graphs display the fluorescence intensity (arbitrary unit) in each channel over the distance depicted by the white arrows. The correlation coefficient ($R^2$) between the fluorescence intensities in the cytoplasm (depicted by the black arrows) is indicated. Scale bar indicates 20 µm. Shown is a representative result from n=4 independent experiments. The histograms display the  the colocalization between the BioCyTASQ and MitoTracker Deep Red (MTdR) signals using the ratio of colocalized areas over total MTdR area (green bars) or the ratio of colocalized areas over total BioCyTASQ area (orange bar). Data are presented as mean values ± SEM of n=4 independent experiments. **(d)** Labeling of HeLa cells with or without BioCyTASQ (treatment of live cells with 1 µM BioCyTASQ for 24 h), with DAPI and treated or not with RNase. Shown is a single representative field from one experiment over n=3 independent experiments. Scale bar indicates 20 µm. **(e)** RG4 detection in fixed cells using the BioCyTASQ probe (1 µM for 24 h) coupled with an immunofluorescence analysis of TOMM20 and IkB protein localization in HeLa cells. Scale bar indicates 10 µm. **(f)** Graphs displaying the fluorescence intensity (arbitrary unit) in each channel over the distance depicted by the red arrow in (e). The correlation coefficient ($R^2$) of the colocalized fluorescence intensities of BioCyTASQ and IkB with TOMM20 signal is indicated and calculated in 5 different cells. Data are provided as a Source Data file. **(g)** Quantification of the BioCyTASQ signal intensity from Supplementary Fig. 1d plotted relatively to the DMSO condition. Data are presented as mean values ± SEM of n=4 independent experiments and each dot is a cell, ***P<0.001 (two-sided paired t-test).

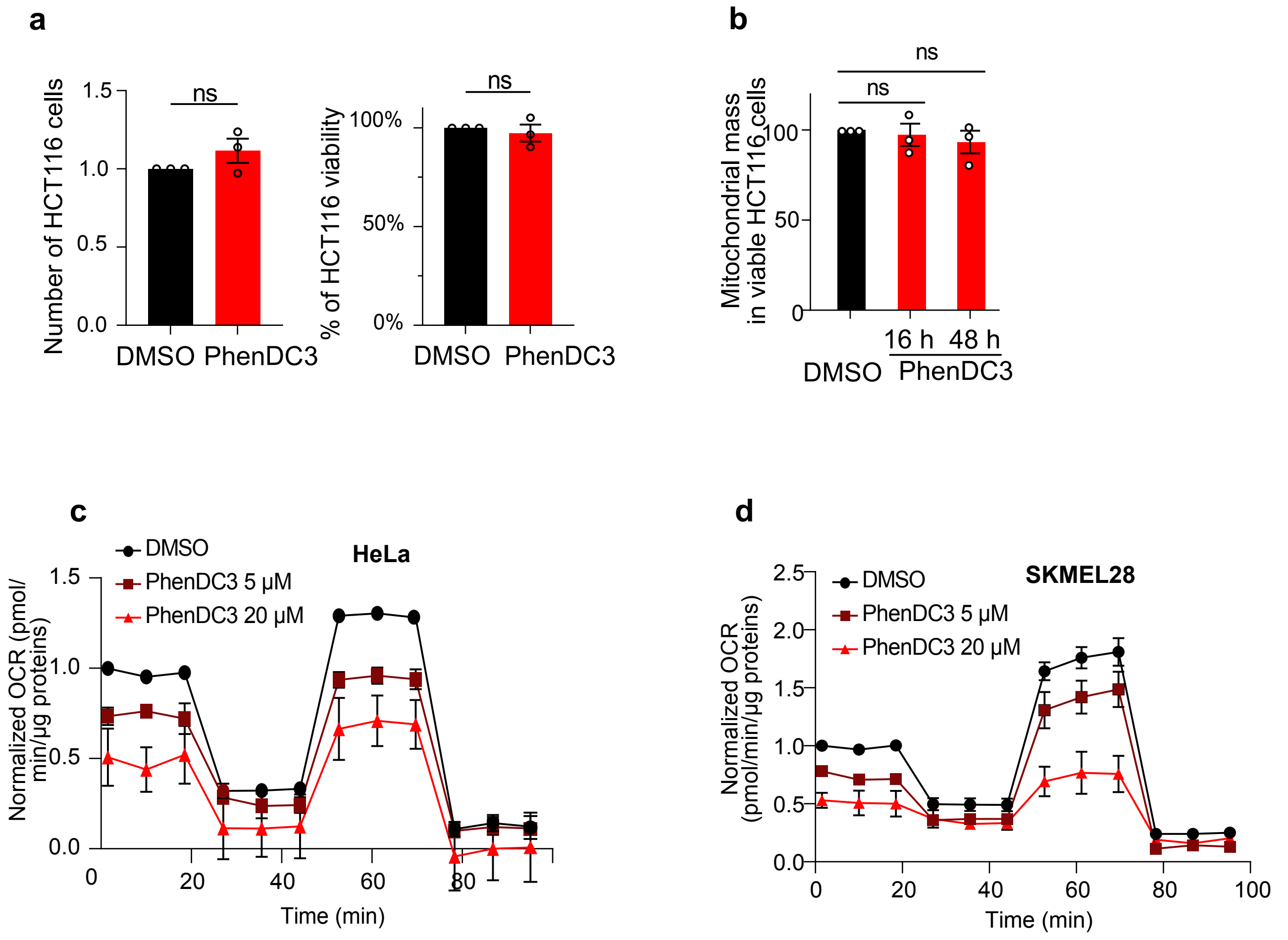


**Supplementary Figure 2 (related to Figure 1). RG4 function in mitochondrial respiration. (a)** HCT116 cell number and percentage of viability, followed by trypan blue cell counting, after treatments with DMSO and 20 μM of PhenDC3 for 24 h. Data are presented as mean values ± SEM of n=3 independent experiments, ns: non-significant (two-sided paired t-test). **(b)** Quantification of the mitochondrial mass followed by flow cytometry analysis of the fluorescent MTG probe of HCT116 cells treated with DMSO or 20 μM of PhenDC3 for 16 h and 48 h and plotted relatively to the DMSO condition. Data are presented as mean values ± SEM of n=3 independent experiments, ns: non-significant (two-sided paired t-test). **(c,d)** Quantification of the normalized Oxygen Consumption Rate (OCR) per cell, measured by a Seahorse assay, in HeLa (c) and SKMEL28 (d) cells treated with DMSO or a dose scale of PhenDC3 for 16 h. Data are presented as mean values ± SEM of n=3 independent experiments. Data are provided as a Source Data file.

Dumas, *et al.*, Supplementary Fig 3

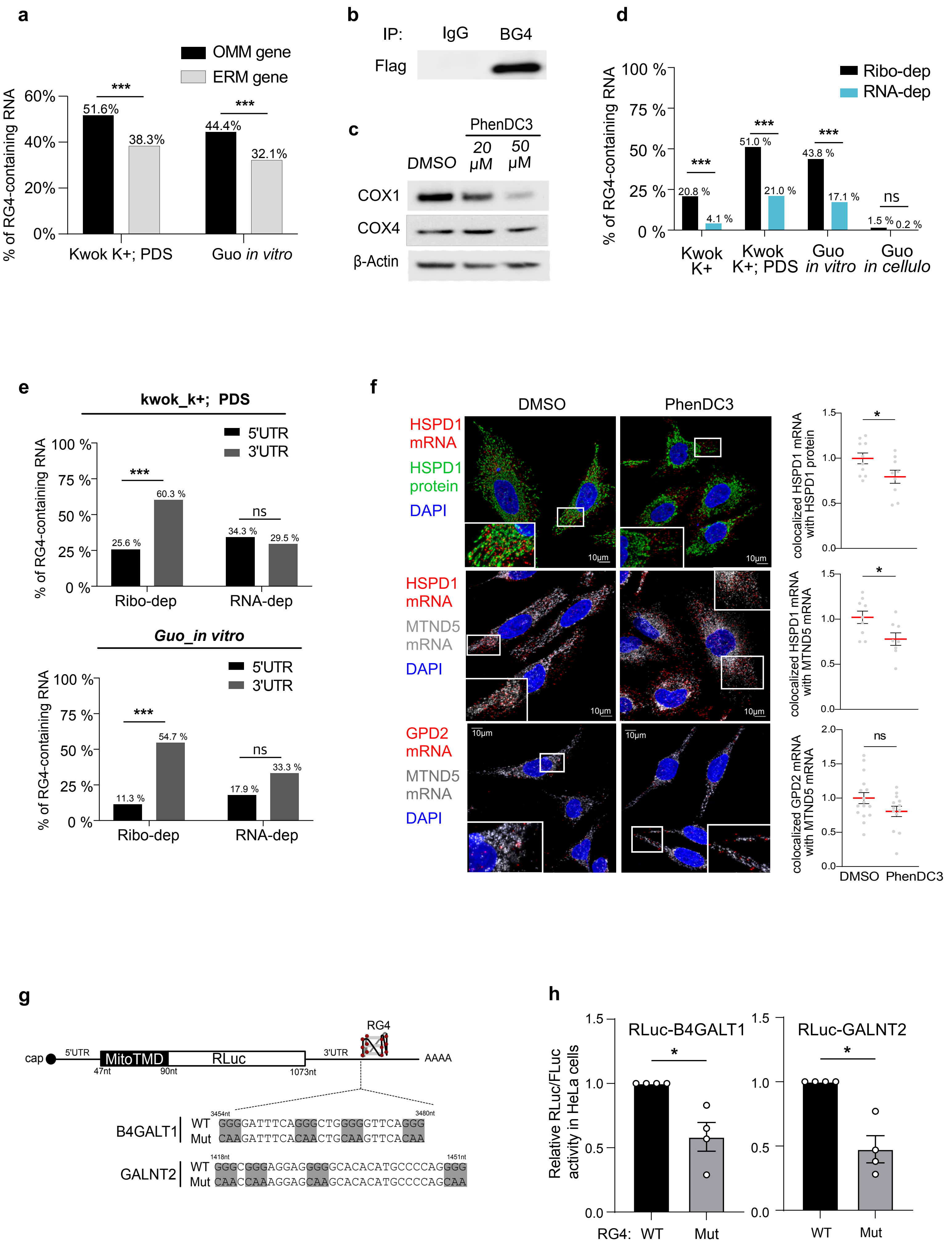

**Supplementary Figure 3 (related to Figure 2). Analysis of RG4s in nuclear-encoded mitochondrial mRNAs (a)** Proportion of RG4-containing mRNAs experimentally validated based on Kwok *et al* and Guo *et* al [1,2] localized at the OMM and the Endoplasmic Reticulum Membrane (ERM) based on Fazal *et al* [4]. For the RG4-containing mRNAs experimentally validated in K+ and cPDS+ conditions based on Kwok *et al,* P=1.297e-09 and for the RG4-containing mRNAs experimentally validated *in vitro* based on Guo *et al*, P = 9.346e-09 (chi square test of proportions OMM vs ERM). **(b)** Immunoprecipitation of *in cellulo* RNA-protein complexes (RIP) in HCT116 cells (cytoplasmic fraction) using the Flag-tagged BG4 antibody or control IgG, followed by western blot analysis of the Flag epitope tag. Shown is a representative result from n=3 independent experiments. **(c)** Western blot analysis of COX1, COX4 and □-Actin in HCT116 cells treated with dose scale of PhenDC3 for 48 h. The blot shown is a representative result from n=3 independent experiments. **(d)** Proportion of RG4-containing mRNAs experimentally validated based on Kwok *et al* and Guo *et* al [1,2] in ribosome (ribo)- and RNA-dependent nuclear-encoded mitochondrial mRNAs based on Fazal *et al* [4]. For the RG4-containing mRNAs experimentally validated in K+ and/or cPDS+ conditions based on Kwok *et al* [2] or in vitro based on Guo *et al* , P<2.2e-16; for the RG4-containing mRNAs experimentally validated in cellulo based on Guo *et al* [1] P=0.07348 (chi square test of proportions ribo- vs RNA-dependent mRNAs). **(e)** Proportion of ribosome (ribo)- and RNA-dependent nuclear-encoded mitochondrial mRNAs based on Fazal *et al* [4] containing RG4 in 5'UTR and 3'UTR based on Kwok *et al* and Guo *et* al [1,2]. For the RG4-containing mRNAs experimentally validated in K+ and cPDS+ conditions based on Kwok *et al*, P< 2.2e-16 and P = 0.5537 for ribo- and RNA-dependent mRNAs, respectively. For the RG4-containing mRNAs experimentally validated in vitro based on Guo *et al, P*< 2.2e-16 and P = 0.5537 for ribo- and RNA-dependent mRNAs, respectively (chi square test of proportions 5'UTR- vs 3'UTR). **(f)** RNAscope analysis of HSPD1, GPD2 and MTND5 mRNA localization and immunofluorescence analysis of HSPD1 protein localization in HeLa cells treated with DMSO, 20 µM PhenDC3 for 24 h. Scale bar indicates 10 µm. Shown is a single representative field from one experiment over n=2 or n=1 independent experiments for HSPD1 or GPD2, respectively. The graphs display the normalized HSPD1 protein fluorescence intensities around the HSPD1 mRNA or the co-localization between HSPD1 and MTND5 or GPD2 and MTND5 and were plotted relatively to the DMSO condition. Data are presented as mean values ± SEM of n=2 or n=1 independent experiments for HSPD1 or GPD2, respectively and each dot is a cell, *P<0.05, ns: non-significant (two-sided paired t-test). Data are provided as a Source Data file. **(g)** Illustration of the capped and polyadenylated mRNA reporters containing the Mitochondrial Transmembrane Domain (MitoTMD) of AKAP1 fused to the Renilla Luciferase (RLuc) open reading frame and containing the 3'UTR of B4GALT1 or of GALNT2. Each 3'UTR regions carry the experimentally identified G-quadruplex forming sequence wild type (WT) or mutated (Mut), in which Gs are replaced by As in each stretch of guanines (in gray) forming the RG4. **(h)** Ratio of Renilla/Firefly luciferase activities (Rluc/Fluc) determined in HeLa cells transfected with RLuc-B4GALT1 and RLuc-GALNT2 reporter mRNAs (capped and polyadenylated depicted in Supplementary Fig. 3g) containing the RG4 unmodified (WT) or mutated (Mut) and an internal control mRNA encoding the Firefly luciferase (Fluc). Data are presented as mean values ± SEM of n=4 independent experiments *P<0.05 (two-sided paired t-test). Data are provided as a Source Data file.

Dumas, *et al.*, Supplementary Fig 4

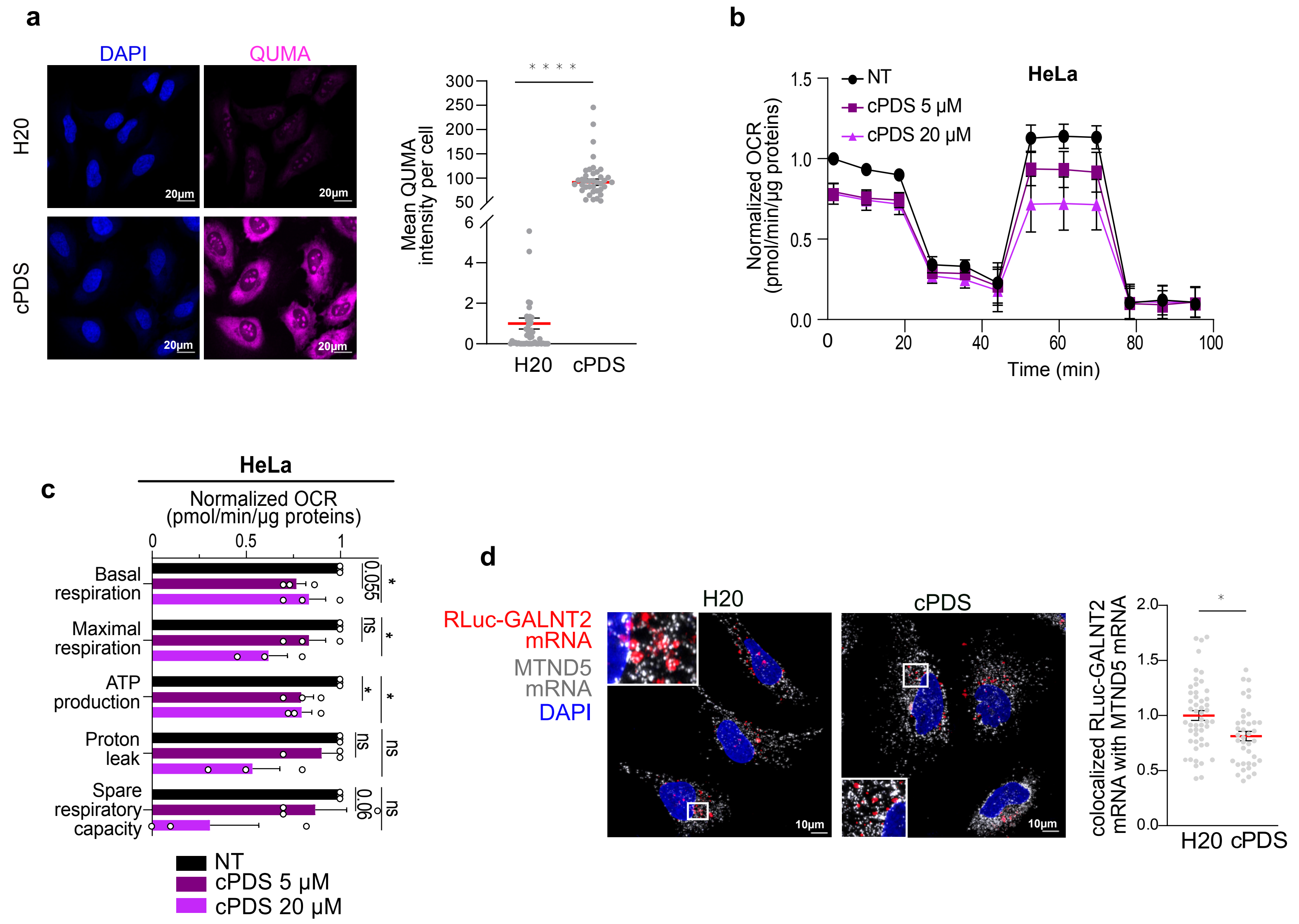


**Supplementary Figure 4 (related to Figure 2). Effect of cPDS on RG4 formation, mitochondrial respiration and RG4-mRNA localization at mitochondria (a)** RG4 detection in fixed cells using the QUMA probe (1 µM for 24 h) in HeLa cells treated with 20 µM of cPDS for 24 h. Scale bar indicates 20 µm. The graphs display the quantification of the QUMA signal intensity per cells plotted relatively to the control condition (H20). Number of cells counted : 49 cells for H20, 43 cells for cPDS. Data are presented as mean values ± SEM and each dot is a cell, **P<0.001, ***P<0.001 (two-sided paired t-test). **(b)** Quantification of the normalized Oxygen Consumption Rate (OCR) per cell, measured by a Seahorse assay in HeLa cells treated with a dose scale carboxypyridostatin (cPDS) for 16 h. Data are presented as mean values ± SEM of n=3 independent experiments. **(c)** Quantification of the Oxygen Consumption Rate (OCR) linked to basal and maximal mitochondrial respiration, ATP turnover, proton leak, and spare respiratory capacity, measured by Seahorse assay, in HeLa cells treated with a dose scale of cPDS for 16 h and plotted relatively to the untreated condition (NT). Data are presented as mean values ± SEM of n=3 independent experiments, *P<0.05, ns: non-significant (two-sided paired t-test). **(d)** RNAscope analysis of Rluc-GALNT2 reporter mRNA (capped and polyadenylated depicted in Supplementary Fig. 3g) and MTND5 mRNA localization in HeLa cells treated with H20 or 20 µM carboxypyridostatin (cPDS) for 24 h. Scale bar indicates 10 µm. Shown is a single representative field from one experiment over n=3 independent experiments. Quantification of the co-localization between RLuc-GALNT2 mRNA reporter and MTND5 mRNA. Data are presented as mean values ± SEM of n=3 independent experiments and each dot is a cell, *P<0.005 (two-sided paired t-test). For all the panels, data are provided as a Source Data file.

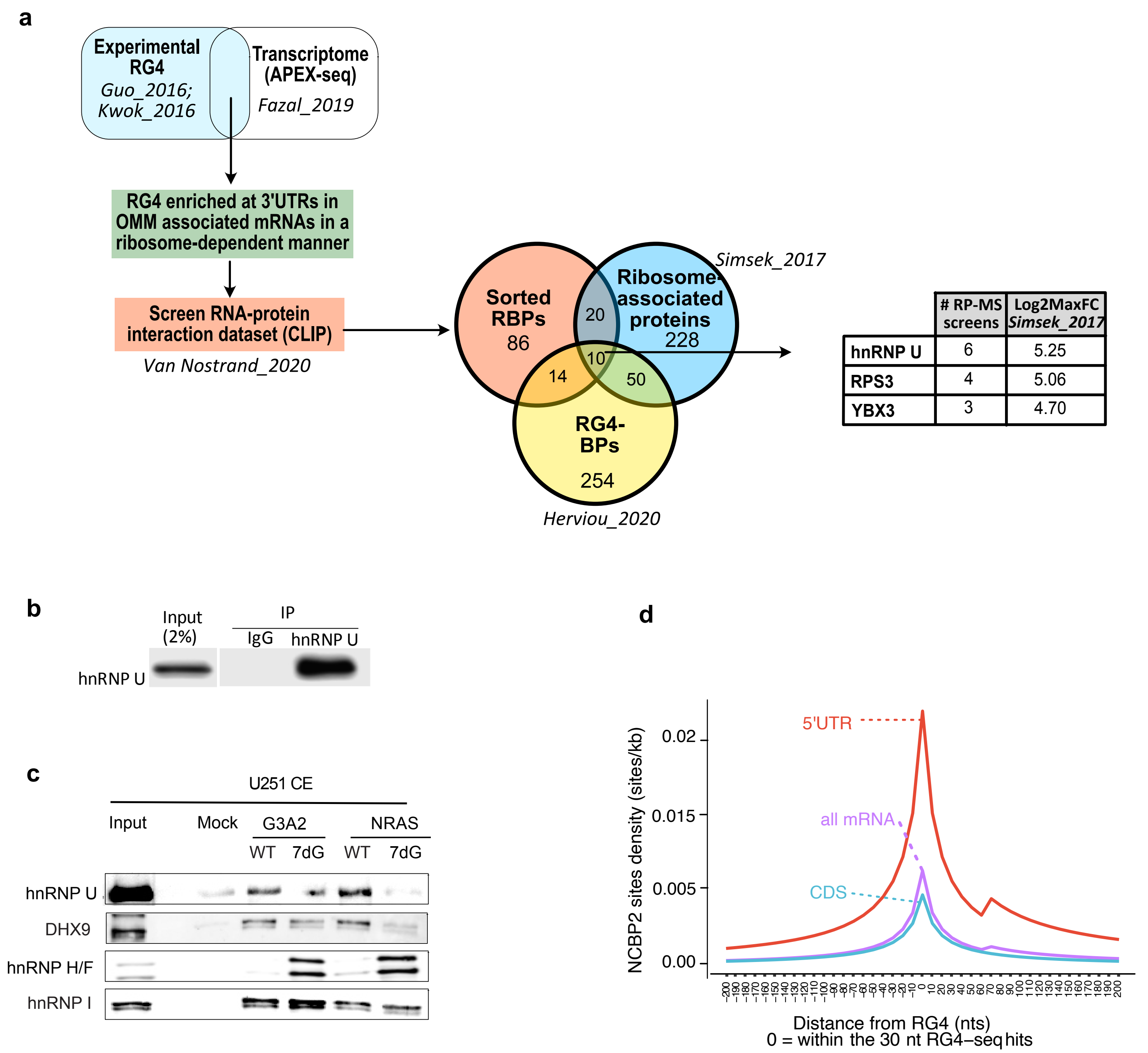


**Supplementary Figure 5 (related to Figure 3). Identification of hnRNP U as a regulator of RG4-containing mitochondrial mRNAs. (a)** Overview of filtering parameters to identify RBPs bound to RG4s located in the 3'UTRs of OMM-associated mRNAs in a ribosome dependent manner mRNAs. **(b)** IP of *in cellulo* RNA-protein complexes in cytoplasmic extracts from HCT116 cells with the hnRNP U antibody. **(c)** RNA affinity chromatography using the G3A2 and NRAS RG4 sequences either native (WT) or 7-deaza-modified (7dG) and U251 cytoplasmic cell extracts, followed by western blot analysis of hnRNP U, DHX9, hnRNP H/F and PTB. Shown is a representative result from n=2 independent experiments for G3A2 sequences, n=1 for NRAS RG4 sequences. Data are provided as a Source Data file. **(d)** Density of NCBP2 binding sites around RG4s (based on [2]) located in mRNA (all mRNA), CDS, and 3'UTR. At distance 0: P = ns (non-significant).

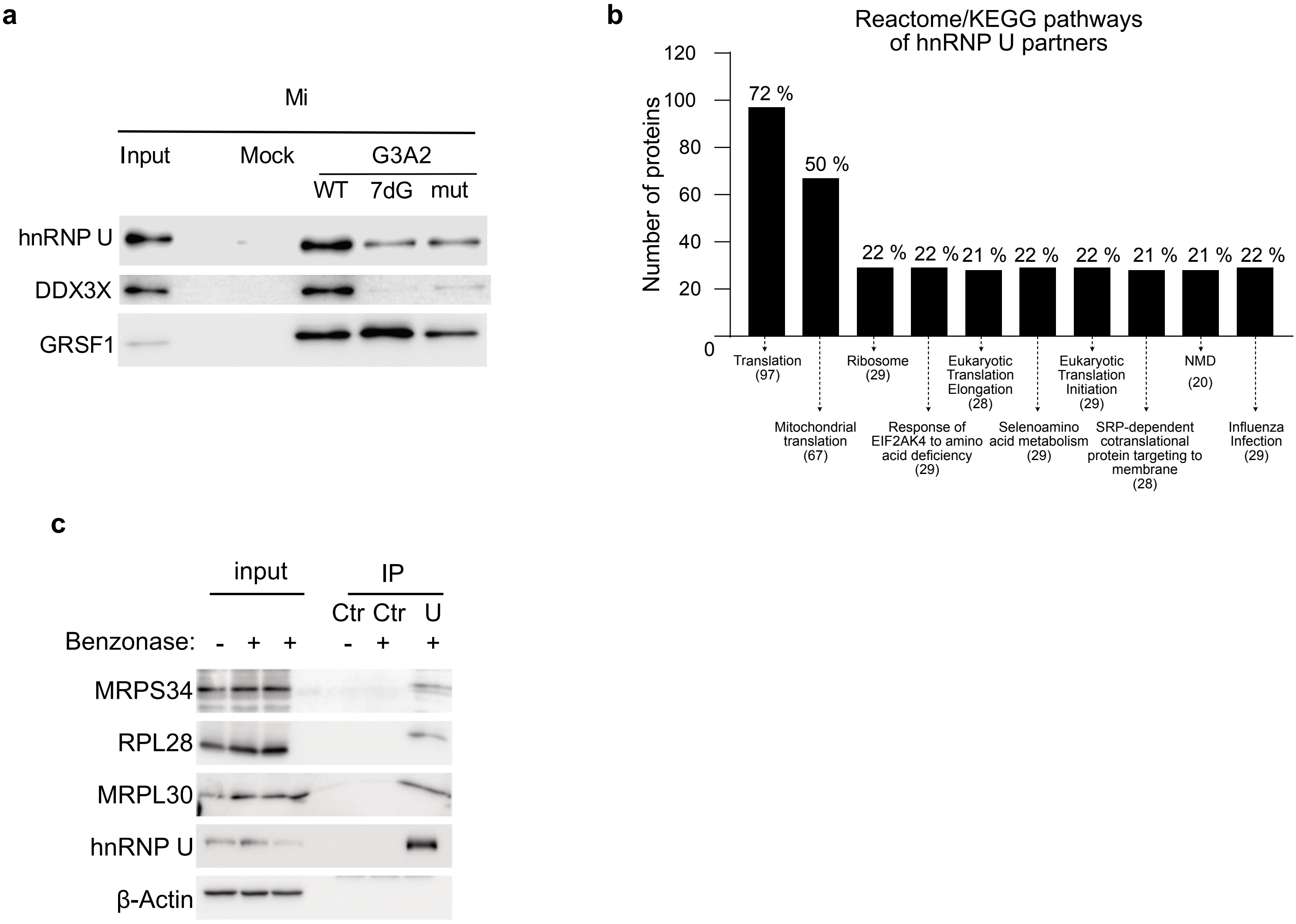


**Supplementary Figure 6 (related to Figure 4). hnRNP U binding to RG4s in mitochondrial fractions (a)** RNA affinity chromatography using the G3A2 and NRAS RG4 sequences either native (WT), 7-deaza-modified (7dG) or mutated (mut) and HCT116 mitochondrial cell extracts (Mi), followed by western blot analysis of hnRNP U, DDX3X and GRSF1. The blot shown is a representative result from n=2 independent experiments. Data are provided as a Source Data file. **(b)** Functional enrichment analysis of the identified high confidence 134 hnRNP U partners. **(c)** Immunoprecipitation (IP) of HCT116 cytoplasmic extracts treated or not with benzonase using the hnRNP U antibody or control IgG, followed by western blot analysis and probing with the indicated antibodies. Shown is a representative result from n=3 independent experiments. Data are provided as a Source Data file.

Dumas, *et al.*, Supplementary Fig 7

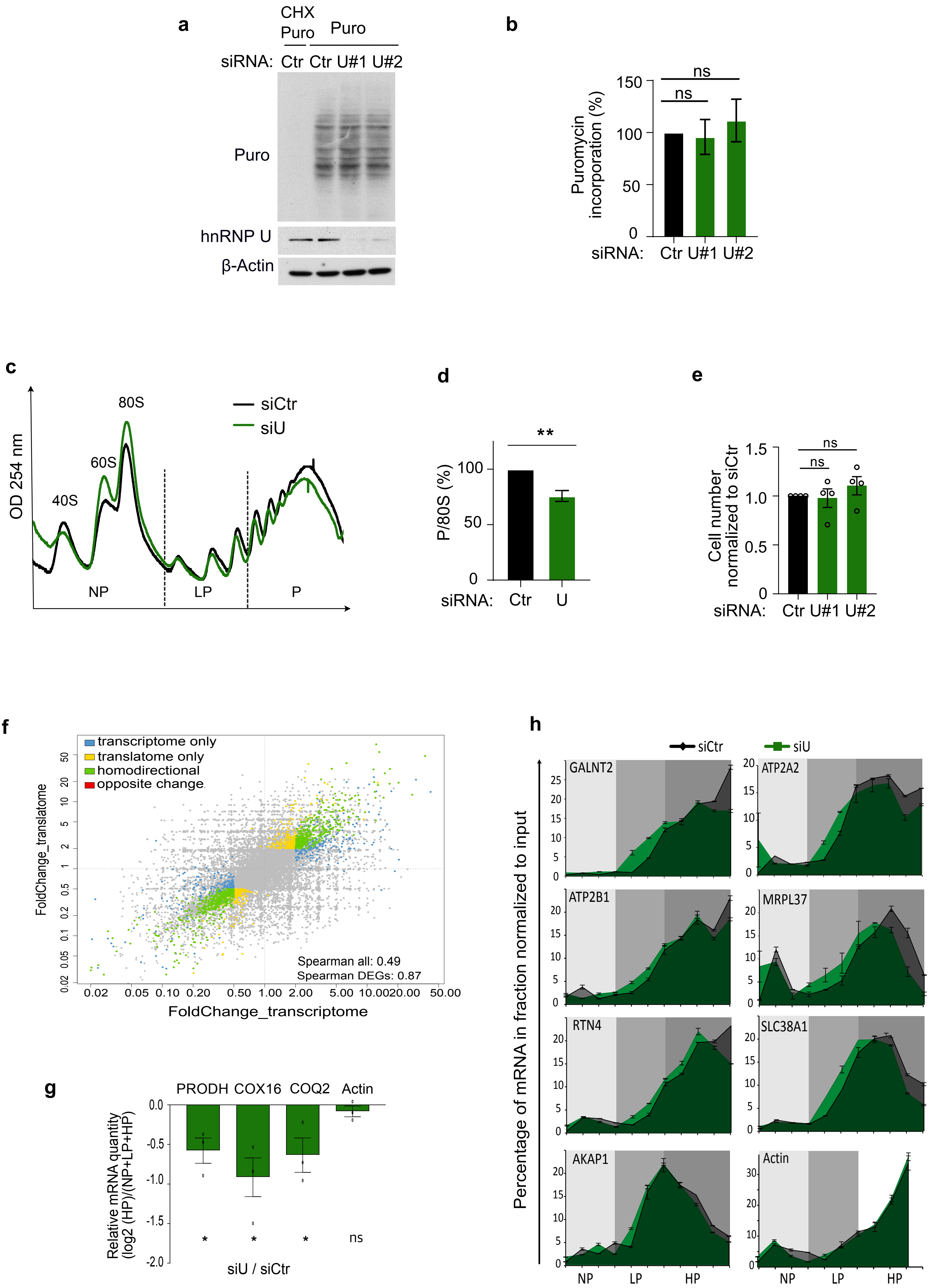

**Supplementary Figure 7 (related to Figure 5). Role of hnRNP U in translational regulation.** **(a)** *De novo* protein synthesis analysis by SUnSET assay in HCT116 cells treated with control (siCtr), and 2 different hnRNP U (siU#1, siU#2) siRNAs, followed by western blot analysis of the incorporated puromycin, hnRNP U and □-Actin. The blot shown is a representative result from n=3 independent experiments. Data are provided as a Source Data file. **(b)** Quantification of puromycin incorporation from (a) plotted relatively to the siCtr condition. Data are presented as mean values ± SEM of n=3 independent experiments, ns: non-significant (two-sided paired t-test). **(c)** Polysome profile of HCT116 cells treated with control (siCtr) and hnRNP U (siU) siRNAs. The positions of the 40S, 60S and 80S ribosomal subunits and non-polysomal (NP) and light (LP) and heavy (HP) polysomal fractions are indicated. Shown is a representative result from n=9 independent experiments. **(d)** Quantification of the ratio between the area under the HP curve and the 80S area from the polysome profile in (c) in the control (siCtr) and hnRNP U-depleted (siU) conditions and plotted relatively to the siCtr condition. Data are presented as mean values ± SEM of n=9 independent experiments, **P<0.01 (two-sided paired t-test). **(e)** HCT116 cell number after treatments with control (siCtr) and 2 different hnRNP U (siU#1, siU#2) siRNAs plotted relatively to the siCtr condition. Data are presented as mean values ± SEM of n=4 independent experiments, ns: non-significant (two-sided paired t-test). **(f)** Transcriptional and translational genome-wide analysis of polysome associated mRNA extracted after polysome profiling of (c). Data are presented as fold change in transcriptome and translatome between HCT116 cells treated with control and hnRNPU siRNAs. Colour-coded are genes with the adjusted P-value < 0.05 with log2(fold change) >=1 (DESeq2); n=3 independent experiment per group. **(g,h)** RT-qPCR analysis from pooled (g) or individual (h) non-polysomal (NP), light (LP) and heavy (HP) polysomal fractions extracted from the polysome profile in (c) using specific primers for the indicated mRNAs. Quantification by analyzing the ratio HP/total mRNAs in (g). Data are presented as mean values ± SEM of duplicates of n=3 independent experiments analyzed in (g) and of a representative experiment from n=2 independent experiments in (h). Data are provided as a Source Data file.

Dumas, *et al.*, Supplementary Fig 8

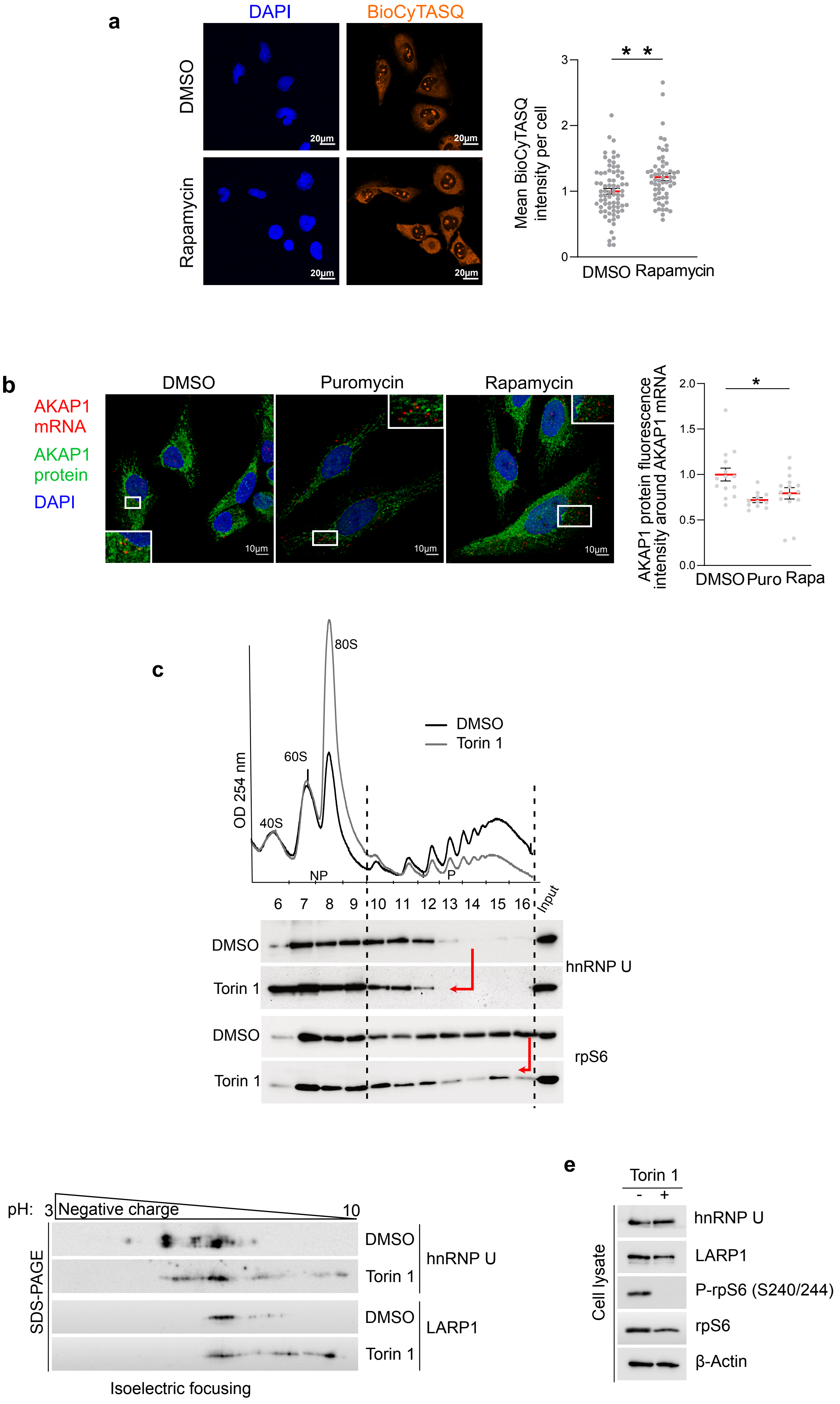

**Supplementary Figure 8 (related to Figure 5). Role of mTOR in hnRNPU-mediated regulation of mitochondrial mRNA translation. (a)** RG4 detection fixed cells using the BioCyTASQ probe (1 µM for 24 h) coupled with an immunofluorescence analysis of TOMM20 protein localization in HeLa cells treated with DMSO or 100 nM of Rapamycin for 2 h. Scale bar indicates 20 µm. The graph displays the quantification of the BioCyTASQ signal intensity plotted relatively to the DMSO condition. Data are presented as mean values ± SEM and each dot is a cell, **P<0.001 (two-sided paired t-test). Number of cells counted: 74 cells for DMSO, 59 cells for Rapamycin. Data are provided as a Source Data file. **(b)** RNAscope analysis of AKAP1 mRNA localization and immunofluorescence analysis of AKAP1 protein localization in HeLa cells treated with DMSO or 100 nM of Rapamycin for 2 h and 100 µg/mL puromycin for 1 h. Scale bar indicates 10 µm. Shown is a single representative field from one experiment over n=3 independent experiments. The graphs display the normalized AKAP1 protein fluorescence intensities around the AKAP1 mRNA and were plotted relatively to the DMSO condition. Data are presented as mean values ± SEM and each dot is a cell, *P<0.05 (two-sided paired t-test). Number of cells counted: 14 cells for DMSO, 11 cells for Puromycin and 15 cells for Rapamycin. Data are provided as a Source Data file. **(c)** Polysome profile of HCT116 cells treated with DMSO or 300 nM Torin 1 for 3 h, followed by western blot analysis from individual non-polysomal (NP) and polysomal (P) fractions by probing for hnRNP U and rpS6 (positive control). Shown is a representative result from n=3 independent experiments. **(d)** Isoelectric focusing followed by western blot analysis of hnRNP U and LARP1 in HCT116 cells treated with DMSO and 300 nM Torin 1 for 3 h. The blot shown is a representative result from n=2 independent experiments. **(e)** Western blot analysis of hnRNP U, LARP1, P-rpS6 (positive control), rpS6 and β-Actin in HCT116 cells treated with DMSO and 300 nM Torin 1 for 3 h. The blot shown is a representative result from n=2 independent experiments.

Dumas, *et al.*, Supplementary Fig 9

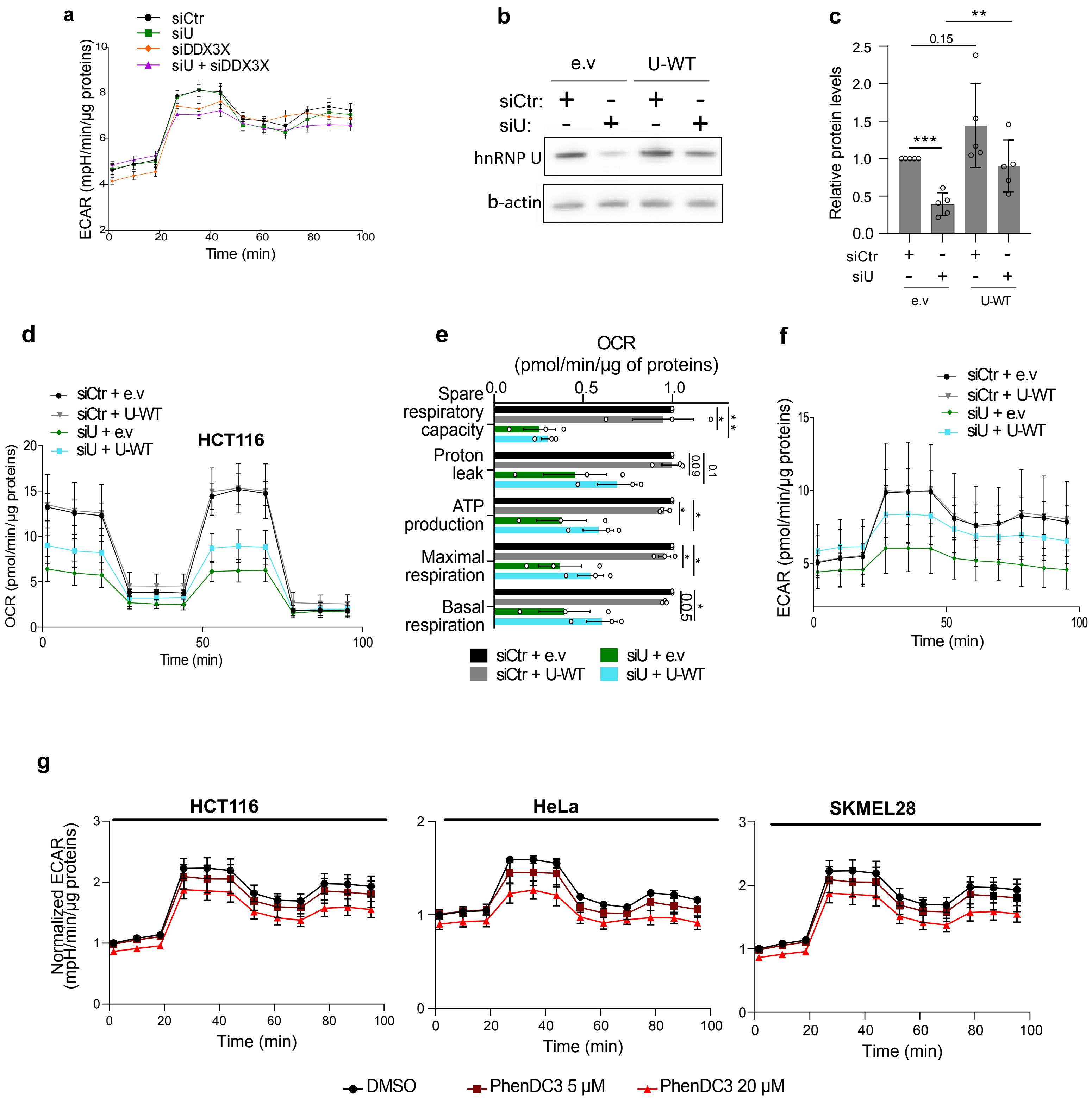


**Supplementary Figure 9 (related to Figure 6). Role of hnRNP U in energy metabolism. (a)** Quantification of the Extracellular Acidification Rate (ECAR) measured by a Seahorse assay (mitochondrial stress test), in HCT116 cells treated with control (siCtr), hnRNP U (siU), DDX3X (siDDX3X) or a combination of hnRNP U and DDX3X (siU + siDDX3X) siRNAs for 48 h. Data are presented as mean values ± SEM of n=3 independent experiments. **(b)** Western blot analysis of hnRNP U and □-Actin in HCT116 cells treated with control (siCtr) or hnRNP U (siU) siRNAs for 72 h and transfected with an empty vector (e.v) or a plasmid expressing hnRNP U WT -FLAG for 48 h. The blot shown is a representative result from n=3 independent experiments.

**(c)** Quantification of the protein levels in (b) normalized to □-Actin and plotted relatively to siCtr condition. Data are presented as mean values ± SEM of n=3 independent experiments, *P<0.05, **P<0.01, ***P<0.001, ns: non-significant (two-sided paired t-test). **(d)** Quantification of the normalized Oxygen Consumption Rate (OCR) per cell, measured by a Seahorse assay in HCT116 cells treated as in (b). Data are presented as mean values ± SEM of n=3 independent experiments. **(e)** Quantification of the Oxygen Consumption Rate (OCR) linked to basal and maximal mitochondrial respiration, ATP turnover, proton leak, and spare respiratory capacity, measured by Seahorse assay, in HCT116 cells treated as in (b). Data are presented as mean values ± SEM of n=3 independent experiments, *P<0.05, ns: non-significant (two-sided paired t-test). **(f)** Quantification of the Extracellular Acidification Rate (ECAR) measured by a Seahorse assay in HCT116 treated as in (b). **(g)** Quantification of the Extracellular Acidification Rate (ECAR) measured by a Seahorse assay, in HCT116, HeLa and SKMEL28 cells treated with with DMSO or a dose scale of PhenDC3 for 16 h. Data are presented as mean values ± SEM of n=3 independent experiments. For all the panels, data are provided as a Source Data file.

Dumas, *et al.*, Supplementary Fig 10

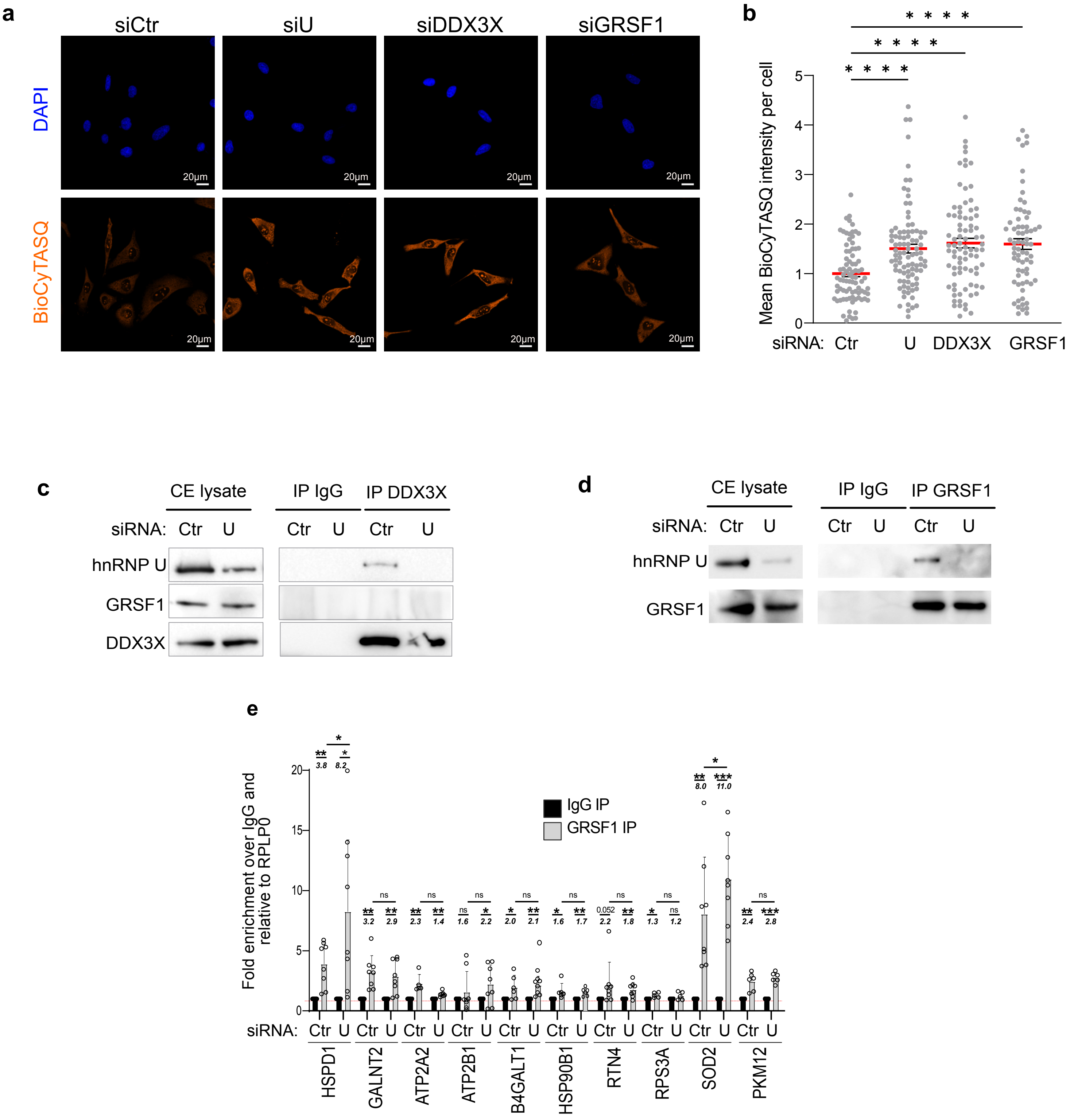


**Supplementary Figure 10 (related to Figure 7). Collaboration between hnRNP U, DDX3X and GRSF1 to regulate RG4-dependent translation. . (a)** RG4 detection using the BioCyTASQ of HeLa cells treated with control (siCtr), hnRNP U (siU), DDX3X (siDDX3X), GRSF1 (siGRSF1) siRNAs for 48 h. Scale bar indicates 20 µm. Shown is a representative result from n=3 independent experiments. **(b)** Quantification of the BioCyTASQ signal intensity from (a) plotted relatively to the siCtr condition. Data are presented as mean values ± SEM of n=3 independent experiments and each dot is a cell, ***P<0.0001 (two-sided unpaired t-test). Data are provided as a Source Data file. **(c)** Immunoprecipitation (IP) with the DDX3X antibody of *in cellulo* RNA-protein complexes in cytoplasmic extracts (CE) from HCT116 cells treated with the control (Ctr) or hnRNPU (U) siRNAs followed by western blot analysis and probing with the indicated antibodies.

**(d)** Immunoprecipitation (IP) of *in cellulo* RNA-protein complexes in cytoplasmic extracts (CE) from HCT116 cells with the GRSF1 antibody. **(e)** RT-qPCR analysis of the indicated mRNAs present in the *in cellulo* RNA-protein complexes from (d). The relative mRNA levels for each IP sample were normalized to the RPLP0 mRNA (negative control) and to the corresponding IP IgG. The italic numbers indicated above each histograms correspond to the fold increase of the mRNA levels in the IP relatively to the IgG. Data are presented as mean values ± SEM of n=3 independent experiments each consisting of duplicates, *P<0.05, **P<0.01, ***P<0.001, ns: non-significant (one-sided paired t-test). Data are provided as a Source Data file.

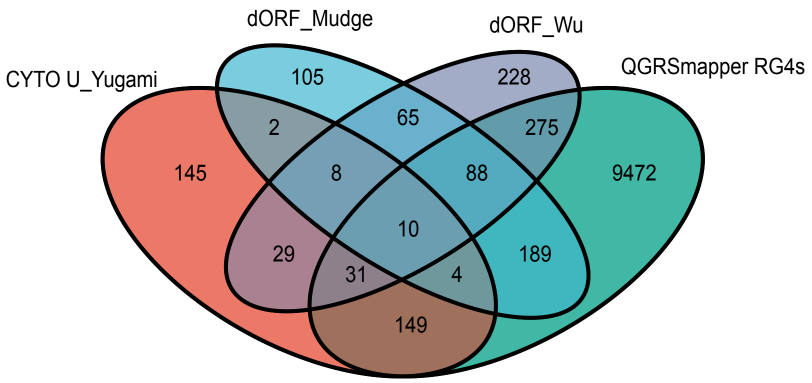


**Supplementary Figure 11. Molecular mechanism underlying translational regulation by 3'UTR RG4s.**Venn diagram showing the overlap of mRNAs Cliped by hnRNP U in the cytoplasm from Yugami *et al*. [5] with the mRNAs containing dORF identified in Wu *et al.*[6] and Mudge *et al.*[7] , and mRNAs containing RG4s predicted by QGRSmapper[8].

Dumas, *et al.*, Supplementary Fig 12

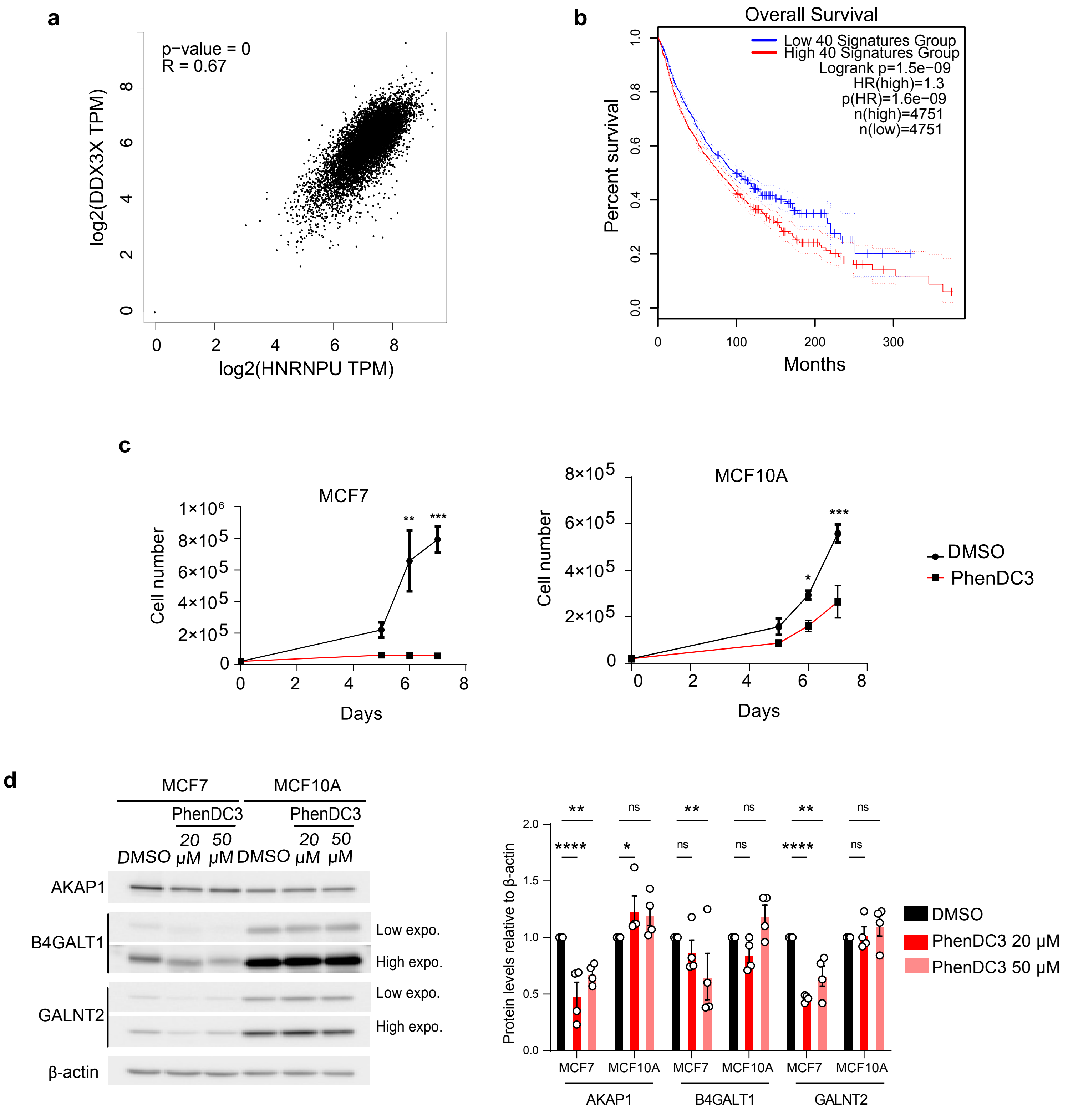


**Supplementary Figure 12. hnRNP U expression and prognostic value in cancer (a)** Expression correlation analysis of hnRNP U and DDX3X using the GEPIA2 database[84]. The Spearman correlation coefficient was used. **(b)** Representation of Kaplan-Meier Overall Survival curve using the GEPIA2 database[84]. **(c)** Proliferation in MCF7 and MCF10A cells treated with DMSO or 20 µM PhenDC3 for the indicated time. Data are presented as mean values ± SEM of n=3 independent experiments. For all the panels, *P<0.05, **P<0.01, ***P<0.001, ns: non-significant (two-way ANOVA). Data are provided as a Source Data file. **(d)** Western blot analysis and quantification of AKAP1, GALNT2, B4GALT1 and [illegible]Actin in MCF7 or MCF10A cells treated with DMSO, 20 or 50 µM PhenDC3 for 48 h. Shown is a representative blot of n=4 independent experiments. Data are provided as a Source Data file.

**SUPPLEMENTARY TABLES**

**Supplementary Table 2.** List of primers used in RT-qPCR and for cloning.

| Primer | Sequence |
|---|---|
| **HPRT_Fw** | TGCTTTCCTTGGTCAGGCAGT |
| **HPRT_Rv** | CTTCGTGGGGTCCTTTTCACC |
| **GPD2_Fw** | ACAACTTGAAACAGCCAGGA |
| **GPD2_Rv** | TCTCTAATACACGCTGAACATCA |
| **ALDHB1_Fw** | AACCCAAGCGTGATCCTGAA |
| **ALDHB1_Rv** | GCTGGTTGTAGGGGATGTCT |
| **PC_Fw** | GGAGAAGGAGCTGGTAGACC |
| **PC_Rv** | CTCCAGCTCCACCTCAAACT |
| **SND1_Fw** | AGGTGCCCCAAGATGATGAT |
| **SND1_Rv** | GAACTGTTTCTCCTTGCGCA |
| **HSPD1_Fw** | AGACAGAGTTACAGATGCCCT |
| **HSPD1_Rv** | TCCTTCAACACCTGCATTCT |
| **GALNT2_Fw** | GGGAACTAACTGCCTCGACA |
| **GALNT2_Rv** | TCTGCTGTCATTTTCTCGGC |
| **PTPRS_Fw** | CCCGCTACCAGTACTTTGTG |
| **PTPRS_Rv** | ACTTGGCCAATGAAGTCGATG |
| **TMEM63B_Fw** | CTCTACTACGCCTACCTGCC |
| **TMEM63B_Rv** | AGACGTGGCAGAGACAGATG |
| **ITM2C_Fw** | TCCCTGGACAAGTGCTATGT |
| **ITM2C_Rv** | CGTTGCACAGGTGGTAGATG |
| **SOD2_Fw** | GACAAACCTCAGCCCTAACG |
| **SOD2_Rv** | CCTGATTTGGACAAGCAGCA |
| **AKAP1_Fw** | CTTCCTCTGATTCAGCTGTGG |
| **AKAP1_Rv** | GTCTACCCACTGGGCAAGG |
| **B4GALT1_Fw** | TGACCATAATGCGTACAGGTG |
| **B4GALT1_Rv** | CTCCCGACCACAGCATTTG |
| **MRPS36_Fw** | CATGATGGGCAGCAAGATGG |
| **MRPS36_Rv** | GAAGAGTGAGATGGTAGCCCT |
| **COX7A2_Fw** | TCTTCGTCAGATTGGGCAGA |
| **COX7A2_Rv** | GTGGCTCTATACAGGAGGGC |
| **USP1_Fw** | ACAGTCCTTAATCATTTCGGTTGA |
| **USP1_Rv** | GGAGTTGGCATGTTTCTTGAATGT |
| **ATP2A2_Fw** | TATGTCGAACCCTTGCCACT |
| **ATP2A2_Rv** | GGCTGCACACACTCTTTACC |
| **ATP2B1_Fw** | ACCCACTGAGTCTCTCTTGC |
| **ATP2B1_Rv** | TCTGAAGGAGGAGCATGCAA |

**Supplementary Table 2.** List of primers used in RT-qPCR and for cloning.

| Primer | Sequence |
|---|---|
| **HSP90B1_Fw** | TGGATCTTGCTGTGGTTTTGT |
| **HSP90B1_Rv** | TGCTCTGTGTCTTCTGTTGTG |
| **RTN4_Fw** | GAACTCAGGCGCCTCTTCTT |
| **RTN4_Rv** | CTATCTGTGCCTGATGCCGT |
| **SLC38A1_Fw** | GGTGATCTTCATACCCTCCATG |
| **SLC38A1_Rv** | AAGGGAATGCTGACCAAGGA |
| **RPS14_Fw** | ATCAAACTCCGGGCCACAGGA |
| **RPS14_Rv** | CTGCTGTCAGAGGGGATGGGG |
| **RPS3A_Fw** | CTTACCCGTGACAAAATGTGT |
| **RPS3A_Rv** | CGTATCTGATTGTTGCGTTT |
| **PKM1/2_Fw** | CATTGCTGTGACCCGGAATC |
| **PKM1/2_Rv** | CCATCCGGTCAGCACAATG |
| **MRPL37_Fw** | GACCTGGACTGTAACGAGGG |
| **MRPL37_Rv** | GCAAACACATTTTGGTCCGC |
| **MRPL53_Fw** | AACGTGGAATCGACGAGGAC |
| **MRPL53_Rv** | AGCATTTCCAGAGCGGTGAG |
| **hnRNP U-FW** | CCGGGATCCGCCGCCACCATGA |
| **hnRNP U-RV-pcDNA3** | CCGGCGGCCGCTTTAGGTGGCGACCGGTGGCTTATCGTCGTCATCCTTGTAATCACGCGTATAATATCCTTGGTGATAATGC |
| **hnRNP U-RV-pEGFP** | CCGGCGGCCGCTTTAGGTGGCGACCGGTGGCTTA |
| **Mut-siRNA-RV** | TGGCGTAGCTCAGCTCGACTTCATCACTTTCAAAGTTAGCAAA |
| **Mut-siRNA-FW** | GTCGAGCTGAGCTACGCCAAGAATGGACAAGATCTTGGCG |
| **B4GALT1-FW** | GCGAGTTCTCAAAAATGAACAATAACGTTTTGGTACACGGATAAGAG |
| **B4GALT1-RV** | TTTTTTTTTTTTTTTTTTTTTTTTTTTTTTTTTTTTTTTTTTTTTTTTTTTTTACAAAGATAGGGTCATTTATTC |
| **B4GALT1- MUT- FW** | CAAGATTTCACAACTGCAAGTTCACAAAGCATTTCCTTTTCCTGGGAG |
| **B4GALT1- MUT- RV** | TTGTGAACTTGCAGTTGTGAAATCTTGAGTAAGGAAACAGGACCCGC |
| **GALNT2-FW** | GCGAGTTCTCAAAAATGAACAATAAGAGGGTCCGGGAGGCCCT |
| **GALNT2-RV** | TTTTTTTTTTTTTTTTTTTTTTTTTTTTTTTTTTTTTTTTTTTTTTTTTTTCTAGAAAGTATCTTCTCTTTATT |
| **GALNT2- MUT- FW** | CAACCAAAGGAGCAAGCACACATGCCCCAGCAAAGCGAGGAGAACTCTTGAAATC |
| **GALNT2- MUT- RV** | TTGCTGGGGCATGTGTGCTTGCTCCTTTGGTTGTGTCCTCAGCTCGCACGCAT |

**Supplementary Table 3.** List of primers for generating the DNA templates used for the synthesis of biotinylated RNAs.

| Primer name | Primer sequence |
|---|---|
| G3A2 WT | tgatacTAATACGACTCACTATAGGtcaacttctactctGGGAAGGGAAGGGAAGGGatcatcatgcatcatctcgctagctaa |
| G3A2 Mut | tgatacTAATACGACTCACTATAGGtcaacttctactctGCGAAGTGAAGTGAAGCGatcatcatgcatcatctcgctagctaa |
| NRAS WT | tgatacTAATACGACTCACTATAGGtcaacttctactctGGGAGGGGCGGGTCTGGGatcatcatgcatcatctcgctagctaa |
| NRAS Mut | TgatacTAATACGACTCACTATAGGtcaacttctactctGCGAGTACCGAGTCTGAGatcatcatgcatcatctcgctagctaat |

Note: Capital letter: T7 promoter; Capital and underlined letter: RG4 sequences; Underlined: NheI restriction site for transcription run-off.
